# Not so scary after all? Decoding the landscape of fear through hormonal responses to risky times and places

**DOI:** 10.1101/2025.05.09.653052

**Authors:** Levi Newediuk, Brett R. Jesmer, Gabriela Mastromonaco, Eric Vander Wal

**Affiliations:** Department of Biological Sciences, University of Manitoba, Canada; Department of Fish and Wildlife Conservation, Virginia Tech, Blacksburg, VA 24060, USA; Reproductive Science Unit, Toronto Zoo, Canada; Department of Biology, Memorial University of Newfoundland, Canada

## Abstract

Prey must balance the energetic benefits of foraging with avoiding predation risk. This risk-reward trade-off, a cornerstone of behavioural ecology, hinges not only on realized predation risk but also on how prey perceive that risk. We often assume energetically rewarding habitats must be inherently risky because prey often increase their vigilance in these habitats or avoid them altogether. However, our assumption that these antipredator behaviours reflect perceived risk frequently goes untested. We used non-behavioural data to test our assumptions about which habitats prey perceive as risky by pairing observations of habitat use of elk (*Cervus canadensis*) with their physiological responses measured from faecal hormones: glucocorticoids (GC), which reflect stress from perceived risk and hunger, and triiodothyronine (T3), which increases with energy intake. Elk had lower GC and T3 in the forest, a putatively safer and poorer foraging habitat than cropland, where they produced more T3, indicating foraging. Surprisingly, GC levels were consistent in cropland, even during the daytime when human activity—and putative risk—peaked. This lack of risk responsiveness highlights that physiological responses are a nuanced integration of perceived risk and reward rather than a guaranteed outcome of habitat use. Our study challenges the assumption that high-reward habitats are inherently risky, and that safer habitats limit energy intake, revealing that the assumptions we make about habitats from a behavioural lens may not always be the reality for prey.

## Introduction

Animals avoid foraging in areas and at times when predators are active (Lima and Dill 1990; Kohl et al. 2019; Smith et al. 2019). For this reason, predators are often required to cue in on habitats that contain the most or best quality food for their prey (Williams and Flaxman 2012). These predator-prey dynamics likely make the most energetically rewarding habitats simultaneously the most dangerous. Consequently, animals are often forced to forage in dangerous places, thereby negatively impacting their survival and reproduction (Creel et al. 2007; Elliott et al. 2017; Gehr et al. 2018). Importantly, it is not just the realized risk of predation that affects animals negatively, but also their perception of risk. Even when perceived risk does not reflect realized risk, perceived risk can still deter individuals from using optimal foraging habitats (Basille et al. 2015), reduce time spent foraging (Yin et al. 2017), and induce chronic stress (Janssens and Stoks 2014). Because of these effects, the spatiotemporal perception of predation risk—known as *the landscape of fear*—has become a foundational idea in behavioural ecology (Laundré et al. 2001). However, where and when we can expect changes in behaviour and associated stress depends on the assumptions we make about the places and times prey perceive risk.

The perception of predation risk in foraging animals is often inferred from their antipredator behaviours. These behaviours include increasing vigilance in risky places (Laundré et al. 2001), or for social animals, joining larger groups to dilute their risk (Hasenjager and Dugatkin 2017). Avoiding risky habitats is also a form of antipredator behaviour (Dwinnell et al. 2019), particularly during times of heightened risk (Smith et al. 2019). However, antipredator behaviours do not always reliably indicate perceived predation risk. For example, animals may avoid risky habitats because competitive interactions restrict access or limit foraging in these often energetically rewarding areas (Vijayan et al. 2012; Dupuch et al. 2014). Conversely, animals may not avoid risky habitats, even when risk is perceived, if territoriality restricts their access to safer habitats (Creel et al. 2023). These scenarios highlight the need for additional, non-behavioural evidence to test assumptions about which places and times are risky and whether accepting those risks is indeed tied to energetic rewards.

Neurochemical signalling hormones, produced when animals perceive risk, could provide empirical evidence to validate long-standing assumptions about what antipredator behavioural responses tell us about habitats (Jesmer et al. 2017). In vertebrates, perceived risk stimulates the hypothalamus to release stress hormones, including glucocorticoids (Sapolsky et al. 2000). While glucocorticoids also respond to diminished energy reserves and fasting (Sapolsky et al. 2000; Ortiz et al. 2001; Laver et al. 2020)and therefore serve as a general stress marker, triiodothyronine—a hormone mainly involved in metabolism—increases when animals forage on high-energy food sources and decreases as energy reserves are depleted (Eales 1988; Dias et al. 2017). Glucocorticoids have been used to assess perceived predation risk across landscapes and periods with different ambient levels of predation (Boonstra et al. 1998; Creel et al. 2009; Le Saout et al. 2016), but are rarely measured in known, free-ranging individuals in ways that link directly to behaviour (but see Newediuk et al. 2024). Together, glucocorticoid and triiodothyronine levels from free-ranging individuals should reveal the risks prey animals actually perceive in habitats assumed to be risky and energetically rewarding (Figure 1).

**Figure 1.**
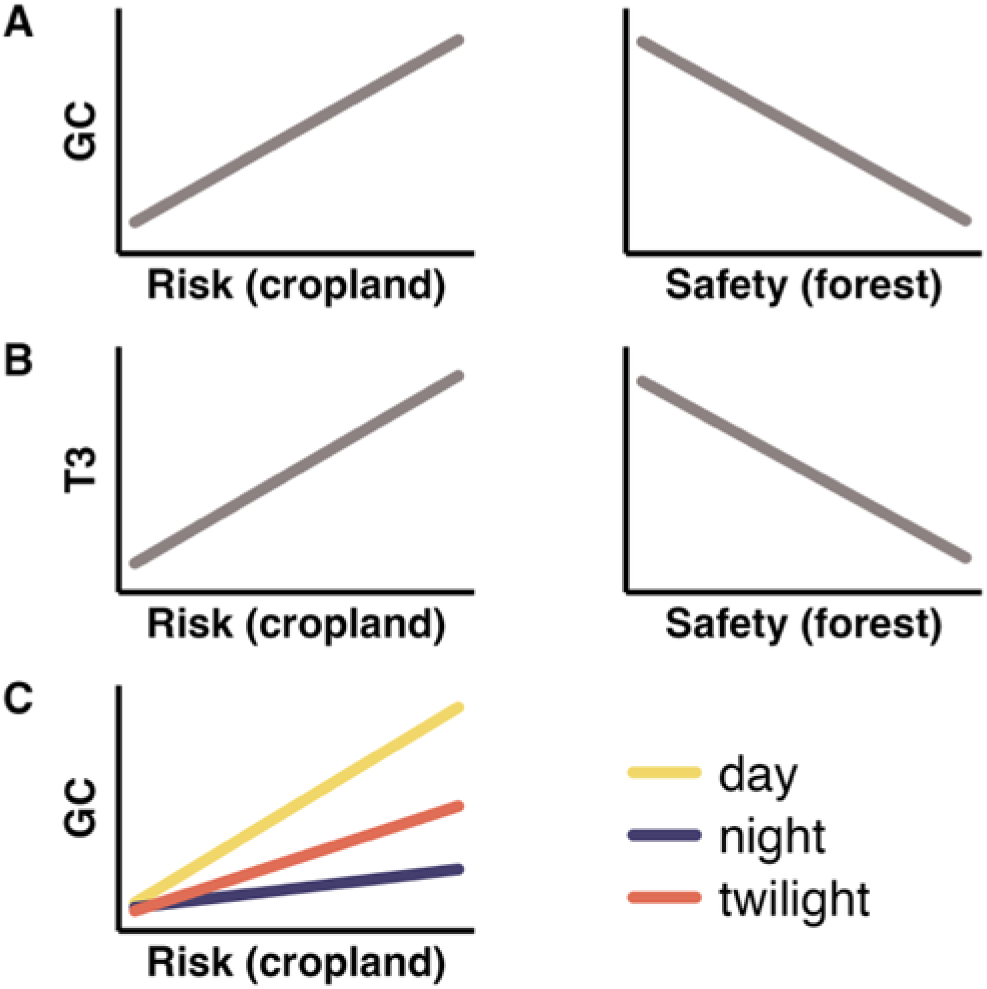
Predicted glucocorticoid (GC) and triiodothyronine (T3) responses in putatively safe and risky habitats. In (A), GC increases after risky habitat use (e.g., cropland) and decreases after use of safe habitat (e.g., forest; Prediction 1), responding to predation risk. In (B), T3 increases after risky habitat use and decreases after safe habitat use (P2), reflecting the assumed link between risk and energetic reward. In (C; P3), GC levels are highest after using risky habitats during risky times (e.g., day) and lowest when predators are inactive (e.g., night and twilight).

We measured glucocorticoid (GC) and triiodothyronine (T3) metabolites to disentangle energetic rewards from the perception of fear by elk (*Cervus canadensis*), a North American ungulate prey species, after using two habitats ostensibly perceived as risky versus safe by the species. Animals are presumed to tolerate risky habitats if potential energetic rewards offset such risk. In contrast, less risky habitats are often assumed to provide less energy. Habitats typically viewed as safe for elk and other ungulates include closed habitats like forests because they provide cover from predators (Amor et al. 2019; Hinton et al. 2020). In contrast, open habitats, such as cropland and other human-altered places, expose ungulates to risk from humans and natural predators (Frid and Dill 2002; Stankowich 2008) but offer substantially greater energetic rewards from crop plants (Brook 2010; Sorensen et al. 2015; DeVore et al. 2016). Since GCs reflect perceived risk (Sapolsky et al. 2000), we predicted GC levels would be highest after elk used cropland—the putatively risky habitat—and lowest after they used forest—the putatively safer habitat (Prediction 1; Figure 1A). Because T3 increases with energy intake (Dias et al. 2017), and animals are assumed to forage in risky places like cropland to access such energy rewards (Brown and Kotler 2004), we also predicted T3 levels to rise after cropland use. Conversely, we predicted T3 to decline after using forest, reflecting less energy intake by elk in this relatively poor foraging habitat (P2; Figure 1B).

Additionally, we tested whether risk perception might vary with time of day. Like many ungulates, elk are crepuscular (Ciuti et al. 2012; Procko et al. 2024), meaning they are most active at dawn and dusk. During the more active crepuscular periods, we expected elk to use cropland for foraging. However, cropland is also a hotspot for human activity, which peaks during the day. The increased human presence should make cropland riskier to elk during the daytime, supported by studies showing that elk and other crepuscular hunted species become increasingly nocturnal with greater human activity in their foraging habitats during the day (Hertel et al. 2016; Visscher et al. 2017). If risk perception changes with the time of day, we predicted elk would exhibit the highest GC levels when they used more cropland during the day, lower levels at dawn and dusk, and the lowest levels at night when human activity is minimal (P3; Figure 1C).

## Methods

### Data collection and processing

We measured summer habitat use by 13 females from a small population of approximately 150 elk in southeast Manitoba, Canada (49.13, -96.56) via GPS collar movements. The collared elk—representing approximately 9% of the population—were captured in February 2019 using a net gun fired from a helicopter and fit with Iridium satellite collars (Vertex Plus 830 g, VECTRONIC Aerospace GmbH, Berlin, Germany) programmed to collect 30-minute GPS locations from May to August 2019 and 2020. Captures were approved in accordance with animal care protocols (Memorial University of Newfoundland AUP #19-01-EV). In summer, our population of elk were most active at twilight and nighttime, foraging in productive cropland interspersed with forest, which they used for resting during the day (Hinton et al. 2020; Figure 2). Forest comprised approximately 40% of the landscape, while cropland covered 12%. The remaining landscape was composed of smaller pockets of wetland and shrubland. Elk primarily used larger blocks of wetland and shrubland surrounding the agricultural core in winter.

**Figure 2.**
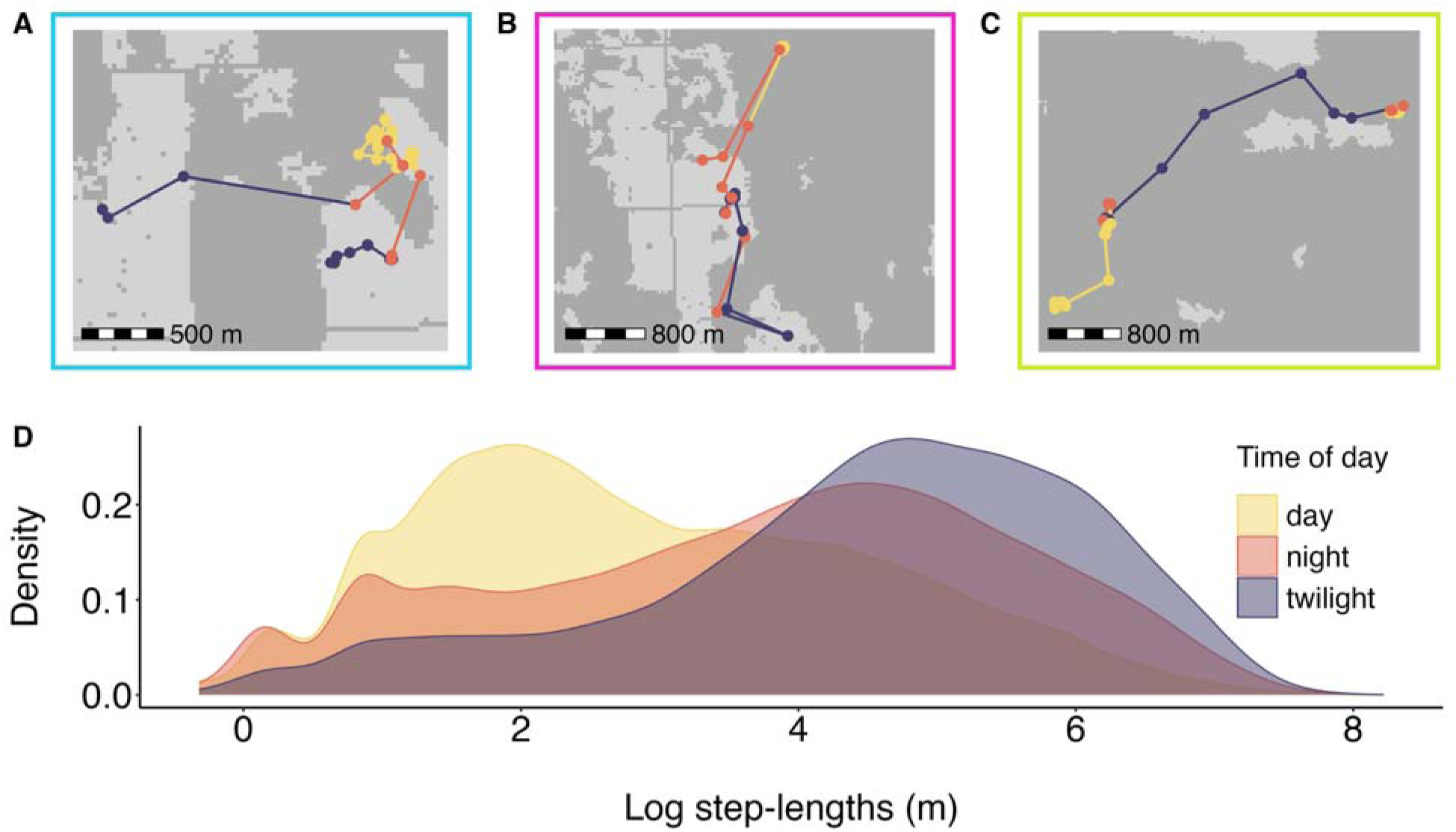
Three examples of elk habitat use strategies in our population, highlighting their use of cropland habitat for foraging rewards while minimizing spatiotemporal risk from human activity. Cropland areas are shown in light grey. Elk locations and tracks, recorded at 30-min intervals for 24 hours (A–C), are coloured according to the time of day: yellow for daytime, orange for twilight, and dark blue for nighttime. The panel border colours (A, blue; B, pink; C, green) correspond to the highlighted points in Figure 3, which shows GC and T3 production by the three elk following the habitat use strategies shown in the panels. Elk that use cropland most extensively forage in open cropland at night and retreat to the forest edge during the day, making short foraging trips into cropland while maintaining proximity to the relative safety of the forest (A). Elk with moderate cropland use restrict their use entirely to nighttime and twilight, avoiding cropland during the day (B). Other elk rarely use cropland, even at night and twilight (C). (D) Illustration of the log-transformed 30-min step length distribution for all 13 collared elk from May to August, indicating they were most active—i.e., they had the longest step lengths—at night and twilight when human activity is minimal.

To quantify GC and T3 levels, we extracted hormones from 75 fresh faecal samples produced by the collared elk, which we collected non-invasively from the landscape over two years in 2019 and 2020. We kept the samples, which we froze no more than 8 hours after their collection, stored at -20° Celsius until we performed the extractions. Our extraction procedure is described in more detail in Newediuk et al. (2024). Briefly, we used an enzyme immunoassay to draw the hormones from dried fecal samples mixed in a methanol-water solution.

We matched each sample to known individuals using nine microsatellites sequenced from the faecal DNA and blood samples collected from the elk at the time of capture. We then assigned a time of defecation to each sample by examining the GPS collar location data to identify when the matched elk was last near the collection site. We used a previously developed machine-learning approach (Newediuk and Vander Wal 2021) to identify any samples lacking sufficient DNA quality for microsatellite sequencing. The machine-learning approach uses movement characteristics and evidence of elk bedding in the area to accurately assign samples to individual elk. Detailed methods are in Newediuk and Vander Wal (2021).

To test the relationship between hormone levels and habitat use, we quantified the proportion of GPS locations in forest and cropland in the hours before defecation. We defined habitat use as the proportional number of times within a set period that an animal’s location was recorded within a land cover type, e.g., forest or cropland, relative to its locations in other habitats during the same period (Lele et al. 2013). We classified elk locations by land cover type using spatial land cover products from Agriculture and Agri-food Canada (AAFC). The AAFC produces annual spatial records of crop cover for agricultural regions in Canada at a spatial resolution of 30 m, based on Landsat-5, AWiFS, DMC and Radarsat-2 satellite imagery. The classification also includes natural land cover types like forests and wetlands. We extracted the land cover type at each elk location point from the corresponding year of spatial imagery—2019 or 2020—using the *sf* package v.1.0-15 (Pebesma 2018) in R v.4.3.1 (R Core Team 2023).

In elk and other mammals, GC and T3 measured in faeces are metabolites of circulating hormones produced by the thyroid and adrenal glands, integrated into faeces over approximately 20–24 hours after the stimulus that caused the production of the hormone (Wasser et al. 2000; Wasser et al. 2010). Our stimuli of interest were the energy intake and risks associated with habitat use. To connect hormonal signals of energy intake and risk with habitat use the previous day, we matched the GC and T3 levels in the samples with the proportional use of forest and crop fields 24 hours prior, beginning 44 hours before the time of the sample to account for the 20 hours required for the hormones to metabolize and become integrated in faeces. We also calculated separate proportions of habitat use during twilight, day, and night in the 24-hour focal period. We distinguished the day, night, and twilight periods by nautical twilight.

### Data analysis

To test the relationship between habitat use and hormone levels, we used Bayesian linear mixed-effects models. We fit two models for each hormone-habitat combination to create four models in total. We fit separate models for forest and cropland to avoid complicating causal interpretations if both habitat types were included in the same model. Because forest and cropland are the most common landcover types on the landscape, and are the two primary habitats used by elk during summer (Hinton et al. 2020), we expected the proportional use of these habitats would be inversely related. Each of the four models included either the proportion of forest use or cropland use as the predictor and faecal T3 or GC levels as the response. We fit a fifth model to test whether the time of day influenced the relationship between cropland use and GC levels. In this model, we included the proportion of cropland use and its interaction with time of day (day, night, or twilight) as predictors, with GC level as the response.

To account for repeat samples from individuals, we included random effects. Although we collected multiple samples from some individuals, most had relatively few samples (median = 5 samples per individual, range = 1–18 samples). We opted for random intercepts to avoid unreliable random slope estimates for the large number of individuals without multiple samples (Wright 2017). We fit all models using the *brms* package v2.20.4 (Bürkner 2017) in R v.4.3.1 (R Core Team 2023). We specified a lognormal error distribution with a log link and used weakly informative priors *N* ∼ 0, 20. We ran all models with four chains and 10,000 iterations, including 5,000 warm-up iterations. We assessed model fit using posterior predictive checks, which compare simulated data from the fitted model to the original data. Agreement between the simulated and observed data indicates a good model fit (Gelman and Hill 2006). We assessed variable significance using the probability of direction, which is the probability that a parameter is entirely positive or negative. Its interpretation is analogous to the frequentist p-value (Makowski et al. 2019).

## Results

We tested the relationship between habitat use, perceived risk, and energetic rewards using 75 faecal samples from 13 elk. Model support was mixed for our habitat use-risk perception predictions. Faecal GC levels did not increase after elk used cropland, a purportedly risky habitat (ß = 0.21, 95% CrI -0.39, 0.81, probability of direction = 75%; Figure 3A), though they did decline after elk used forest, a purportedly safe habitat (ß = -0.33, 95% CrI -0.71, 0.07, p. direction = 95%; Figure 3B). As predicted, faecal T3 levels increased when elk used more cropland in the previous 24 hours (ß = 0.45, 95% Credible Interval -0.23, 1.13, probability of direction = 90%; Figure 3C) and decreased after they used more forest (ß = -0.41, 95% CrI -0.87, 0.06, p. direction = 96%; Figure 3D).

**Figure 3.**
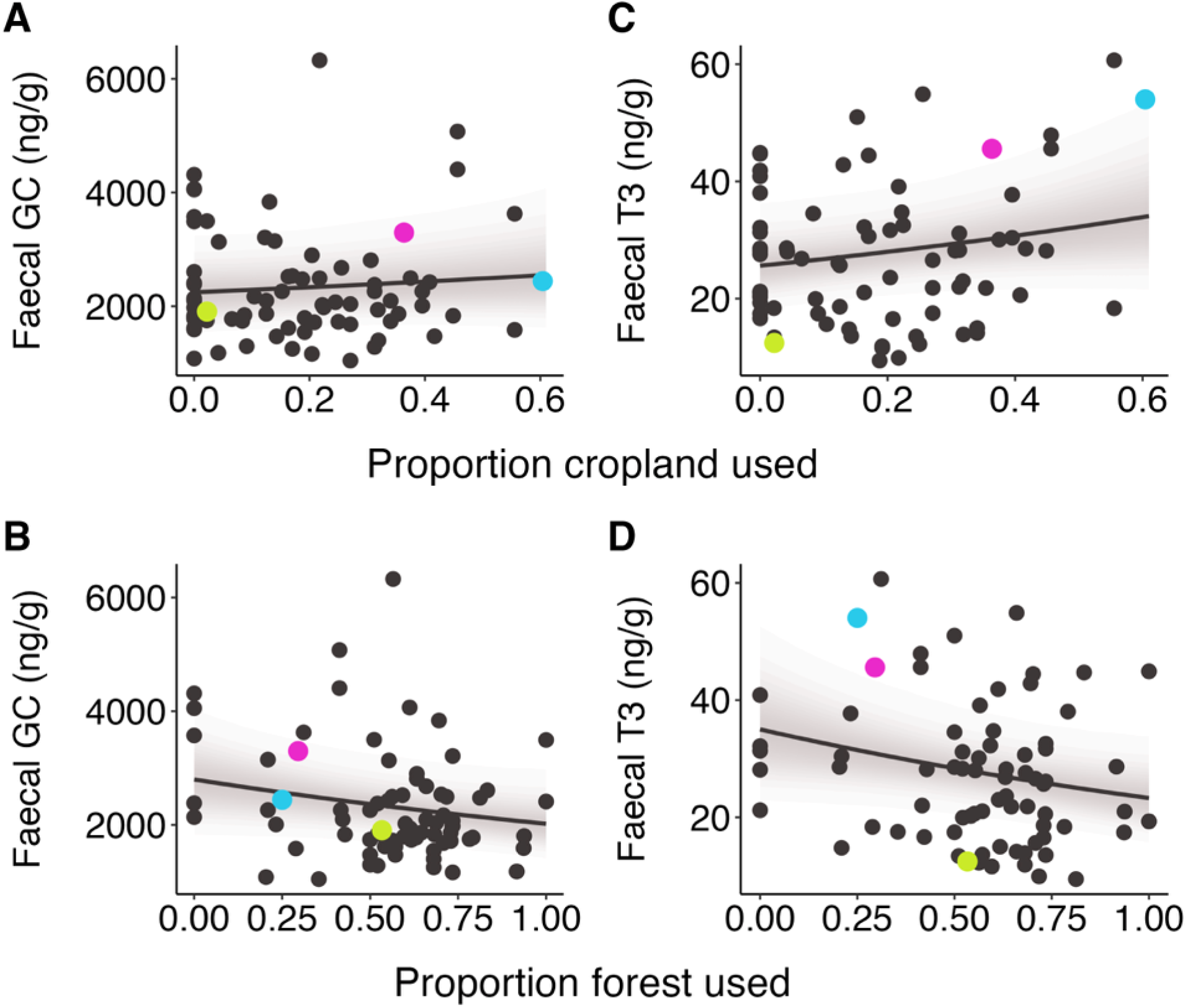
Elk foraged more in cropland than in forest habitats, where they also perceived little risk. However, they appeared not to perceive appreciably more risk in cropland. The panels illustrate changes in faecal hormone metabolite levels—glucocorticoids (GC) and triiodothyronine (T3)—after elk used different proportions of cropland (A, C) and forest (B, D) in the previous 24 hours. The grey bands delineate the 95% credible intervals around the posterior mean (solid line). Black points show the observed faecal GC and T3 levels, reflecting perceived risk and energy intake. The coloured points match the panel borders in Figure 2, which shows three habitat use strategies corresponding to the habitat proportions used here: blue corresponds to foraging in open cropland at nighttime and twilight with short foraging trips into cropland from the safer forest edge by day, pink corresponds to cropland use only at nighttime and twilight, and green corresponds to almost complete avoidance of cropland.

We also tested whether the habitat use-risk perception relationship varied by time of day and found no support for the prediction that faecal GC levels after cropland use would peak during the daytime along with purported risk. The relationship between daytime cropland use and faecal GC levels did not differ from the relationship with twilight (ß = 0.30, 95% CrI -0.57, 1.18, p. direction = 75%; Figure 4) or nighttime cropland use (ß = 0.26, 95% CrI -0.61, 1.11, p. direction = 72%; Figure 4).

**Figure 4.**
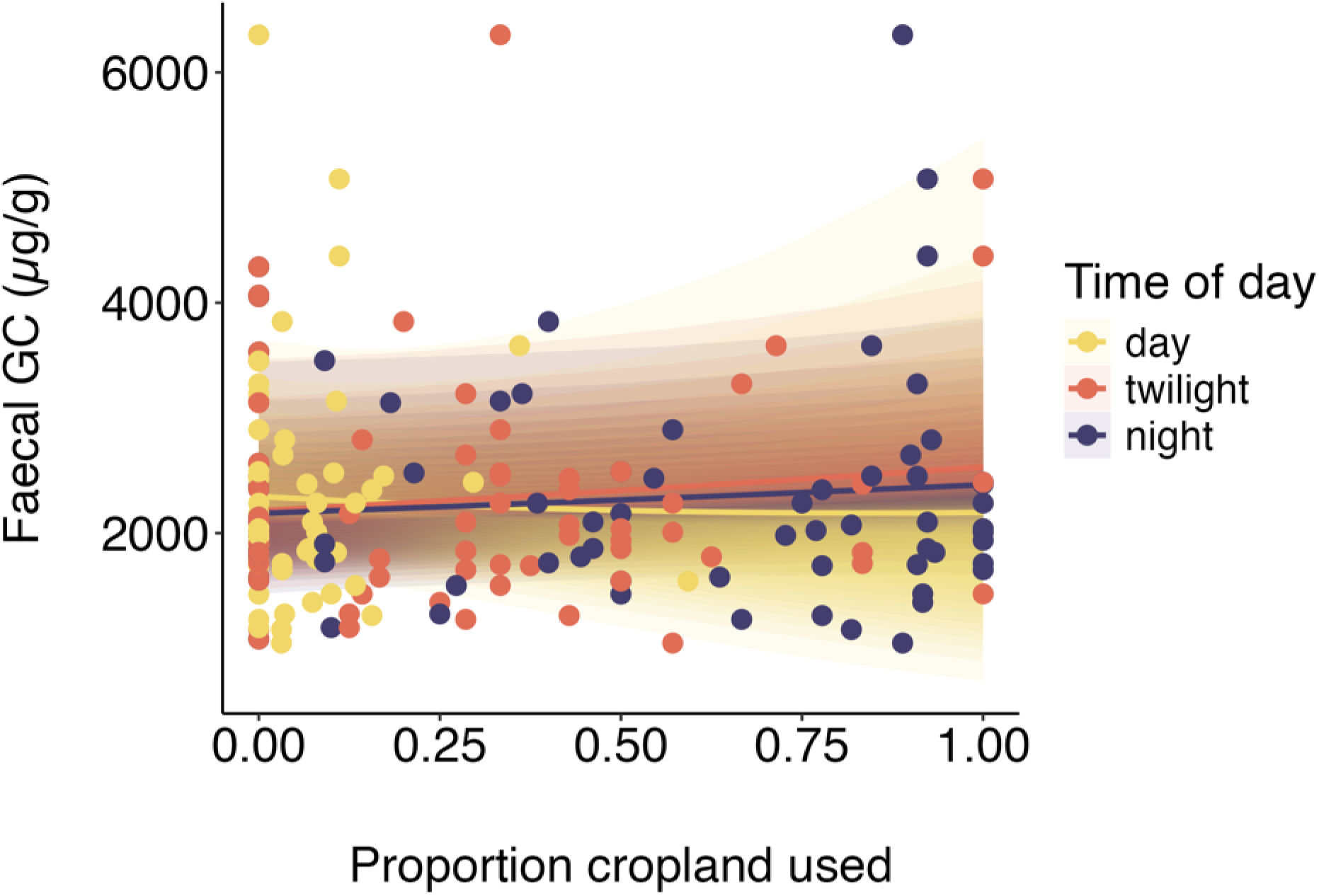
Faecal glucocorticoid (GC) levels, indicating perceived risk, did not change after elk used different proportions of cropland during the day, twilight, and night in the 24 previous hours. The coloured bands delineate the 95% credible intervals around the posterior mean (coloured solid lines). Points show the observed faecal glucocorticoid levels, coloured by time of day.

All models converged successfully (Rhats = 1.0, all ESS ≥ 4,721). Posterior predictive checks indicated that the fitted model, with a lognormal response distribution, fit the observed data well (Supplemental Figures S1 & S2).

## Discussion

Habitat use is strongly influenced by balancing access to energetic rewards with avoiding predation risk (Lima and Dill 1990). To avoid risk, animals reduce their use of energetically rewarding habitats (Basille et al. 2015) and spend less time foraging (Yin et al. 2017). These antipredator behaviours are often interpreted as a trade-off between risk and reward, yet whether animals actually perceive predation risk and sacrifice energetic rewards remains unclear. Using fecal hormone metabolite analysis, we measured the physiological consequences of habitat use. Triiodothyronine (T3) reflects energy intake, and glucocorticoids (GC) indicate psychological stress from perceived risk. As predicted (Prediction 1), we found that elk perceived forests—often regarded in the literature as low-risk habitats for ungulates (Amor et al. 2019; Hinton et al. 2020)—as safe. Greater forest use was indeed followed by lower GC production. T3 levels in faeces indicated that elk aquired less energy in forests and more in cropland (P2), a habitat often considered highly energetically rewarding but risky (Brook 2010; Sorensen et al. 2015; DeVore et al. 2016). Surprisingly, GC levels did not rise after cropland use, contrary to the expectation that energetically rewarding habitats are inherently risky (P1; Brown and Kotler 2004). Moreover, GC levels stayed consistent regardless of when elk used cropland, despite the assumption that risks to ungulates are amplified in cropland in the daytime when humans are active (P3; Paton et al. 2017; Visscher et al. 2017). These findings suggest animals might tolerate risky habitats when the energetic payoff is sufficiently high, provided they leave for safer habitats before the risk outweighs the reward.

GC levels were slightly elevated, but did not increase significantly, as the proportion of time spent in cropland increased. However, GCs declined as elk spent more time in the forest, suggesting elk perceived the forest as a safer habitat relative to cropland and used it to avoid risk. The difference in GC levels between cropland and forest validates earlier findings that elk use forests as refuge habitats during periods and in places that are assumed to be risky (Paton et al. 2017; Hinton et al. 2020). T3 also declined in the forest (Figure 3D), indicating elk were not actively foraging, further cementing this habitat’s primary role as a refuge. However, the concurrent decline in both hormones also suggests that elk resting in the forest did not incur a significant cost in lost foraging opportunities, at least not enough to elevate their stress levels. Since GC also rises during fasting (Ortiz et al. 2001) and when high-quality forage is limited (Laver et al. 2020), we would expect GC to remain high if elk had been forced to forgo foraging in cropland due to risk. Instead, the net decrease in GC in the forest habitat suggests elk could afford to rest there without a substantial energetic cost, challenging the assumption that landscapes of fear universally impose a trade-off between safety and energetic needs. Further supporting the absence of a marked trade-off between risk and reward in our study, some elk spent nearly the entire 24-hour period preceding their hormone sample in the forest (Figure 2C) while also maintaining the lowest GC levels (Figure 3B).

When elk foraged in cropland, the benefits appeared to outweigh the risks. T3 increased in cropland, as expected for animals foraging on high-quality food (Eales 1988; Figure 3A), while GC remained stable (Figure 3C). Failure to detect increasing GC as a function of time spent foraging in cropland is notable, as GC should increase proportionally with perceived risk (Sapolsky et al. 2000). Instead, any stress the elk experienced in cropland was likely brief and transient, and thus we were unable to detect the changes. If brief or transient changes in GC occurred, they were likely integrated over the 24 hours required for the hormones to metabolize and deposit into their faecal samples (Wasser et al. 2000; Wasser et al. 2010). GCs typically spike during acutely stressful events like predator encounters to stimulate the actions required to respond (Sapolsky et al. 2000), but the low levels we measured suggest that most of the time, elk foraging in cropland were not psychologically stressed. Instead, the energetic rewards of foraging in cropland and its moderating effect on GCs likely masked any evidence of risk perception.

If GC does not increase with time spent in their most energetically rewarding habitat, why do elk not spend all their time in that habitat? Elk likely avoided sustained elevated GCs by engaging in habitat use strategies that proactively mitigate their risk while foraging in cropland. One such strategy was to limit their time spent in cropland. No elk spent more than 60% of the 24 hours we measured in cropland (Figure 3), and most also avoided cropland during the day, with individual use during daytime mostly below 30% (Figure 4). When elk did forage extensively in cropland during the day, they tended to remain near the habitat’s edges (e.g., Figure 2A), possibly to facilitate quick returns to the safety of the forest while maintaining their access to high-quality food. Remaining proximal to escape habitat may have helped them keep their GC levels low, as they only ventured farther into open cropland when active at night and during twilight (Figure 2). This ability to respond proactively to predation risk is thought to moderate GC levels in other systems (e.g., Creel et al. 2009; Hammerschlag et al. 2017). Notably, the individual elk highlighted in Figure 2A that used the most cropland of any elk in our sample produced near the average amount of GCs while also producing the most T3 of any elk (Figure 3), suggesting she acquired the most energy while keeping her stress levels stable.

It has long been assumed that animals need to trade off eating with the risk of being eaten, making energetically rewarding habitats inherently risky. Since Lima and Dill (1990) formalized this fundamental idea, it has become a keystone assumption in ecology and has shaped how we collect and interpret behavioural data. We sought to validate this assumption using physiological stress indicators linked to fear and hunger. Our results suggest that at times, animals may indeed perceive the habitats we label risky as such, but this perception is nuanced and tied closely to context. We found that safe habitats are consistently perceived as safe and less energetically rewarding, evident in hormones linked to hunger. However, putatively risky habitats may not trigger a psychological response unless the risk becomes great enough to offset the energetic rewards they offer. Stress hormone levels fluctuated relatively little at the temporal scale we investigated, likely because animals use behavioural strategies to balance these risks and rewards while foraging. Examining habitat use in tandem with hormones—especially those responsive to both risk and reward—offers a powerful way to characterize habitats. Going forward, this dual approach will likely provide the most complete understanding of how animals navigate the landscape of fear (Pritchard et al. 2020).

## Supplementary Materials

**Figure S1.**
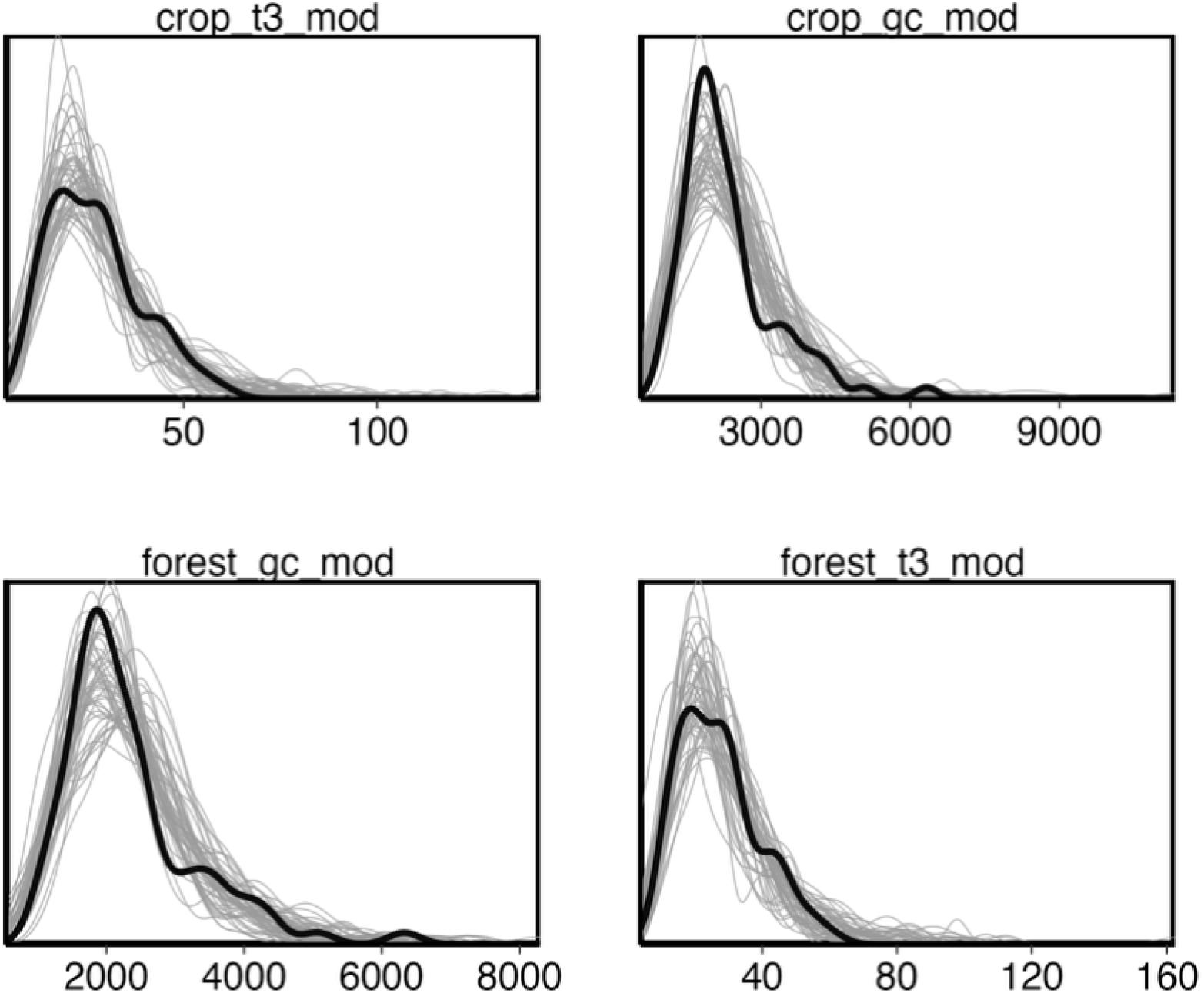
Posterior predictive checks for Bayesian generalized linear models. The distribution of the response variables (bold lines) overlap with simulations from the posterior predictive distributions (spaghetti lines), indicating the models fit the data well. The panels correspond to the models in Figure 3: crop_t3_mod = T3 ∼ cropland use, crop_gc_mod = GC ∼ cropland use, forest_gc_mod = GC ∼ forest use, forest_t3_mod = T3 ∼ forest use.

**Figure S2.**
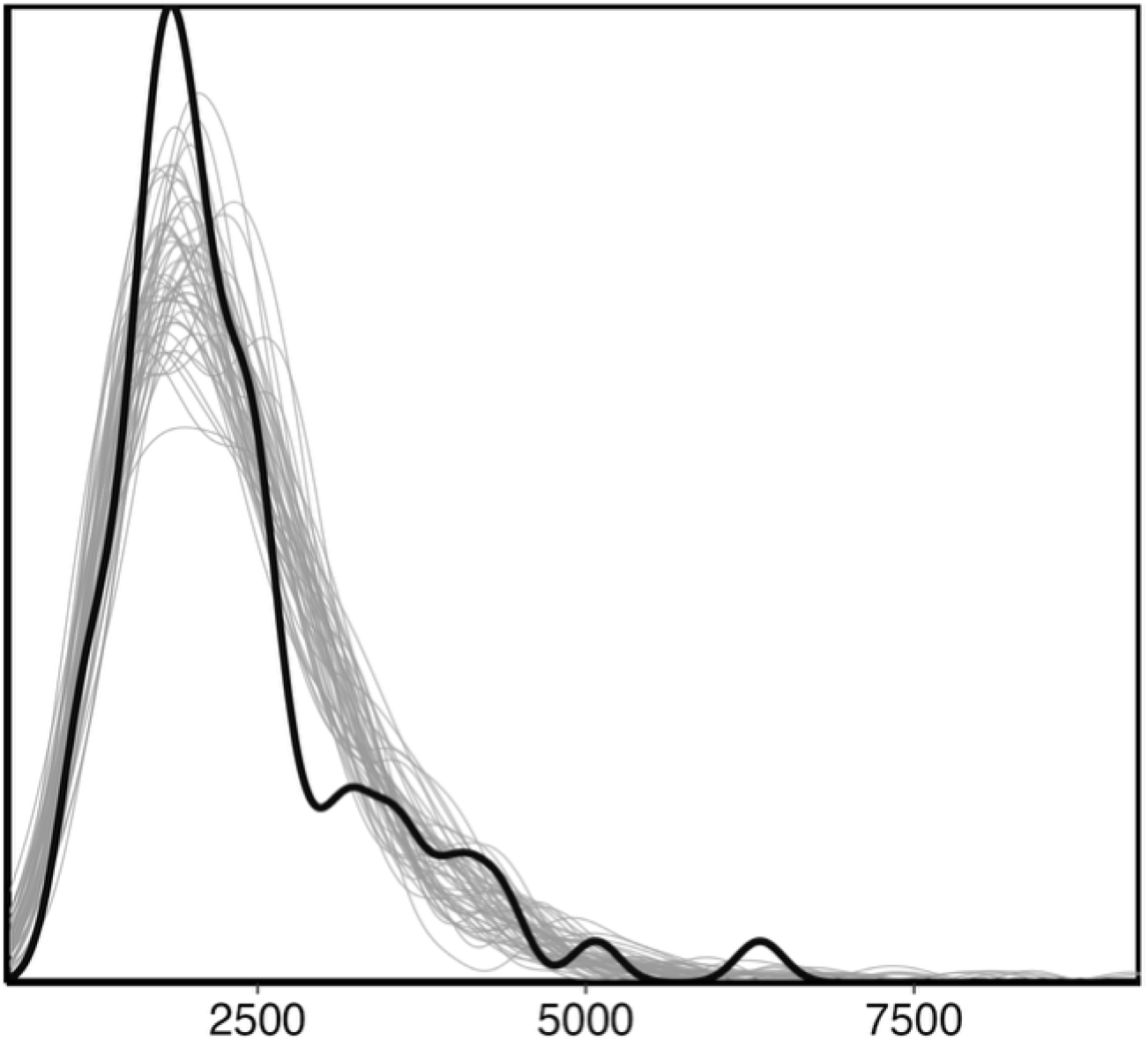
Posterior predictive checks for the Bayesian generalized linear model GC ∼ time of day * cropland use in Figure 4. The distribution of the response variable (bold lines) overlaps with simulations from the posterior predictive distribution (spaghetti lines), indicating the model fits the data well.

